# Restoration of Higher-Order Cortical Processing in Rat V2 by Retinal Sheet Transplants in Degenerated Rats

**DOI:** 10.64898/2025.12.21.690582

**Authors:** Amir-Mohammad Alizadeh, Bin Lin, Magdalene Seiler, David C. Lyon

## Abstract

Age-related macular degeneration (AMD) is a leading cause of central vision loss, and effective treatment options are limited once photoreceptors and the retinal pigment epithelium (RPE) are lost. In advanced stages, vision restoration requires strategies that replace or bypass degenerated retinal circuitry. Retinal sheet transplantation using fetal neural retinal tissue has emerged as a promising intervention, demonstrating long-term survival, integration with the host retina, and partial restoration of light-driven responses. We previously showed that such transplants can restore fundamental visual response properties in the primary visual cortex (V1) of rapidly degenerating rats. However, it remains unclear whether restored retinal input can support higher-order cortical computations that depend on the integration of classical and extra-classical receptive field mechanisms.

In this study, we extend this investigation by evaluating whether fetal retinal sheet transplants can restore extra-classical surround modulation in neurons of higher visual areas (V2). Fetal retinal sheets (E18–E19) derived from donor rats were transplanted into one eye of Rho-S334ter line-3 rats at ages P41–P78, when rod degeneration is nearly complete and cones are largely nonfunctional. Animals were assessed 2.2–9.3 months post-surgery using in vivo extracellular single-unit recordings from V2, optokinetic testing, OCT imaging, and histology. Control groups included normal-vision rats, age-matched degenerated rats (AMC), and sham-operated line-3 rats.

Transplants survived long term, developed laminated and rosetted photoreceptor structures, and integrated with the host retina. Optokinetic testing revealed significant improvement in spatial acuity in transplanted eyes compared with degenerated controls beginning at three months post-surgery. Transplanted rats exhibited a markedly higher proportion of visually responsive V2 neurons than degenerated animals (21.0% vs. 8.2%). They also showed significantly shorter response latencies and larger visually evoked response amplitudes, indicating improved transmission of retinal signals to the cortex.

To quantify surround suppression, neurons were tested with sinusoidal gratings confined to the classical receptive field and gratings extended to full-field size. Transplanted rats displayed robust surround suppression properties similar to normal controls, including significantly reduced firing rates and narrower tuning under full-field conditions. A Support Vector Machine (SVM) classifier trained on net responses to CRF and FF stimulus sizes reliably distinguished control and transplanted neurons from degenerated ones but could not separate control from transplant, further indicating similar response properties in these two groups.

These findings provide the first demonstration that retinal sheet transplants restore not only basic visual responses but also higher-order cortical mechanisms involving extra-classical surround suppression. This recovery of surround suppression in V2 suggests that transplanted retinal tissue can re-establish functionally meaningful circuits capable of supporting complex visual processing. The results underscore the therapeutic potential of retinal sheet transplantation for advanced retinal degenerative disease and provide the first evidence of surround suppression in the rat V2.

## Introduction

Age□related macular degeneration (AMD) is a leading cause of central vision loss in older adults worldwide, with prevalence expected to rise to ∼288 million people by 2040 (Fleckenstein et al., 2021). Macular degeneration is characterized by the accumulation of extracellular deposits (drusen), changes in the retinal pigment epithelium (RPE), choroidal thinning, and photoreceptor degeneration. Cutting edge treatments using micronutrient supplements (Berson et al., 2004) and gene therapy to introduce trophic factors or to correct mutated genes (Bertolotti, Neri, Camparini, Macaluso, & Marigo, 2014; Kauper et al., 2012; Lipinski, Thake, & MacLaren, 2013; M. M. Liu, Tuo, & Chan, 2011; Schwartz et al., 2015; Tsai et al., 2015) can help in the early stages of retinal degeneration where some photoreceptors remain and can therefore be rescued. However, in later stages, once photoreceptors and RPE cells are lost, vision can only be enabled by replacing or bypassing damaged retinal cells. In the more severe and faster degenerating *Rho-S334ter* line-3 rat model (abbreviated as “line 3” throughout the text), and even in the slower degenerating Royal College of Surgeons rat model, cortical responses to visual stimulation are almost entirely abolished in adults (Chen et al., 2016; Coffey et al., 2002; Gias et al., 2011; Girman, Wang, & Lund, 2003), severely limiting normal visual perception. To replace lost photoreceptors in our rodent model of severe retinal degeneration, *Rho-S334ter* line-3 rats, several studies have successfully transplanted fetal retinal sheets (containing neural retinal progenitors) into the subretinal space (Seiler et al., 2014; Seiler, Sagdullaev, Woch, Thomas, & Aramant, 2005; Seiler, Thomas, Chen, Arai, et al., 2008,Seiler, 2017 #75).The fetal retinal sheet transplants survive long-term (Seiler, Aramant, & Ball, 1999), integrate with the host retina (Seiler et al., 2010; Seiler, Thomas, Chen, Wu, et al., 2008), and evoke responses to flashes of light in the superior colliculus, a primary midbrain target of retinal ganglion cells (Sagdullaev, Aramant, Seiler, Woch, & McCall, 2003; Thomas, Aramant, Sadda, & Seiler, 2005; Woch, Aramant, Seiler, Sagdullaev, & McCall, 2001; Yang et al., 2010 ,Seiler, 2017 #75). However it remains unclear how these retinal sheet transplants can restore basic stimulus features, such as orientation tuning, temporal and spatial frequency, contrast, size, and direction which generally elicit very selective responses across visual cortices and are considered key building blocks for the perception of complex shapes and motion (Glickfeld, Reid, & Andermann, 2014; Kobatake & Tanaka, 1994; Livingstone & Hubel, 1988; Marshel, Garrett, Nauhaus, & Callaway, 2011).

We previously compared these response features of V1 cells of the transplanted rats with those of the rats with normal vision and their degenerated age-matched rats. We showed that 3 months after transplantation, the transplanted rats had significantly more visually responsive cells compared to degenerated animals. Also, orientation and direction selectivity as well as other parameters were significantly improved within the parts of V1 which correspond to the location of the transplant (Foik et al., 2018).

Neurons in the primary visual cortex (V1) code basic image features like local orientation, contrast, spatial and temporal properties. Neurons in higher visual areas combine inputs from V1 and code more complex visual properties to eventually build what is known as ‘conscious vision’.

(Coogan & Burkhalter, 1993; Juavinett & Callaway, 2015).

This study furthers our previous study on V1 and assessed the effectiveness of retinal sheet transplants on restoring visual processing, focusing on extra-classical surround suppression in V2 neurons. Using in vivo extracellular recordings, we presented drifting sinusoidal gratings under two conditions: (1) confined to the classical receptive field and (2) extended to full-field size to engage both classical and extra-classical regions. Surround suppression is thought to play a key role in enhancing contrast sensitivity and improving signal-to-noise ratios during visual processing (Y. J. Liu, Hashemi-Nezhad, & Lyon, 2015; Vinje & Gallant, 2000). This phenomenon has been well-documented in primates and cats (Cavanaugh, Bair, & Movshon, 2002; Hashemi-Nezhad & Lyon, 2012; Van den Bergh, Zhang, Arckens, & Chino, 2010), and rodents (Self et al., 2014). By quantifying the response attenuation between the two stimulus conditions, we aim to elucidate the functional role of surround suppression mechanisms in shaping visual representations in rat V2 that received retinal transplant.

To do so, healthy dissected fetal rat retinal sheets were transplanted in *RhoS334ter* line-3 rats at the age of 41-78 days, when degeneration of the rods is nearly complete and cones are largely dysfunctional (Hombrebueno et al., 2010; LaVail et al., 2018; Martinez-Navarrete et al., 2011; Seiler et al., 2014; Zhu, Ji, Lee, & Grzywacz, 2013). Between 2.2 and 9.3 months following transplantation (average 6.9 months), detailed neuronal responses to an array of visual stimuli integral to higher visual processing were measured and compared with control degenerated animals that did not receive transplants, as well as rats with normal vision. These experiments are the first to examine transplant-driven responses and connectivity at the higher cortical level in a rodent model of retinal degeneration and represent an essential step for determining the efficacy of such transplants in visually impaired humans.

## Materials and Methods

### Animals

Animals were treated in accordance with NIH guidelines for the care and use of laboratory animals, the ARVO Statement for the Use of Animals in Ophthalmic and Vision Research, and under two protocols approved by the Institutional Animal Care and Use Committee of UC Irvine.

For this study, 11 non-nude (immunocompetent) rats of the RhoS334ter line 3 strain (*SD-Foxn1 Tg(S334ter)3LavRrrc*; RRRC#539) (8F, 17M) received retinal sheet transplants derived from rat E18-E19 embryos (of these, 4F, 7M were recorded), 13 rats received sham surgery. 29 RD rats (7F, 19M) were used as age-matched non-surgery RD controls and 14 “PA” rats were used as normal retinal controls (8F, 6M).

*Donors for retinal transplantation:* Eight timed-pregnant rats of the ACI-Tg (ROSA26-ALPP) Rrrc strain with normal retina (“PA” rats) were used a source for the donor retinal sheets. These rats originally expressed universally human alkaline phosphatase (hPAP), but this transgene was lost around the time the experiments for the project started. Therefore, we imported breeders of a GFP-expressing line (F344-Tg(UBC-EGFP)F455Rrrc; RRRC#307) and bred them with the existing strain to produce EGFP-expressing “GPA” rats. EGFP tissue was used for two of the 11 cortically recorded transplant rats. To obtain timed-pregnant rats, rats were mated overnight and checked for evidence of mating (sperm check) in vaginal smear. On day 18-20 of gestation (day of conception = day 0), embryos were removed by C-section, and stored in Hibernate E medium (Thermofisher, A12476-01) with B27 supplements (Thermofisher, A3582801) on ice. Retinas were dissected free from surrounding tissues and stored overnight at 4°C in 50-100 µL of Hibernate E medium, sometimes with BDNF/GDNF microspheres (Yang et al., 2010). Tissue was flattened out by capillary force.

*Transplantation procedure*. The transplantation procedure was performed according to previously described methodology (Aramant & Seiler, 2002). Rats received a subcutaneous injection of Ketoprofen (4mg/kg) (Parsippany-Troy Hills, NJ) and dexamethasone eye drops (Bausch & Lomb Inc., Rancho Cucamonga, CA) prior to anesthesia to prevent eyelid swelling. *Line-3* rats (P41 –P78) were anesthetized with a mixture of ketamine (40–55 mg/kg) and xylazine (6 –7.5 mg/kg; i.p.). Pupils were dilated with 1% atropine ophthalmic solution, and local anesthesia was provided by tetracaine eye drops (0.5%) (MWI). A small incision (width 1.0 mm) was made posterior to the pars plana, parallel to the limbus. The implantation instrument was inserted with extreme care to minimize disturbance of the host retinal pigment epithelium. The graft tissue (∼1mm^2^) was released into the subretinal space posteriorly at the nasal quadrant near the optic disc (diagram in **Fig. 1**). Transplants were placed into the left eye only, leaving the right eye as a control. The incision was closed with 10–0 sutures, and the eyes were treated with gentamycin and artificial tears ointment. For recovery, rats were given a subcutaneous injection of Ringer’s saline solution, antisedane (0.25 mg/kg), and the analgesic Buprenex (0.03 mg/kg) for pain management, and placed in a Thermocare incubator. Animals with transplant misplacement and/or excessive surgical trauma were excluded from the study.

**Figure 1.**
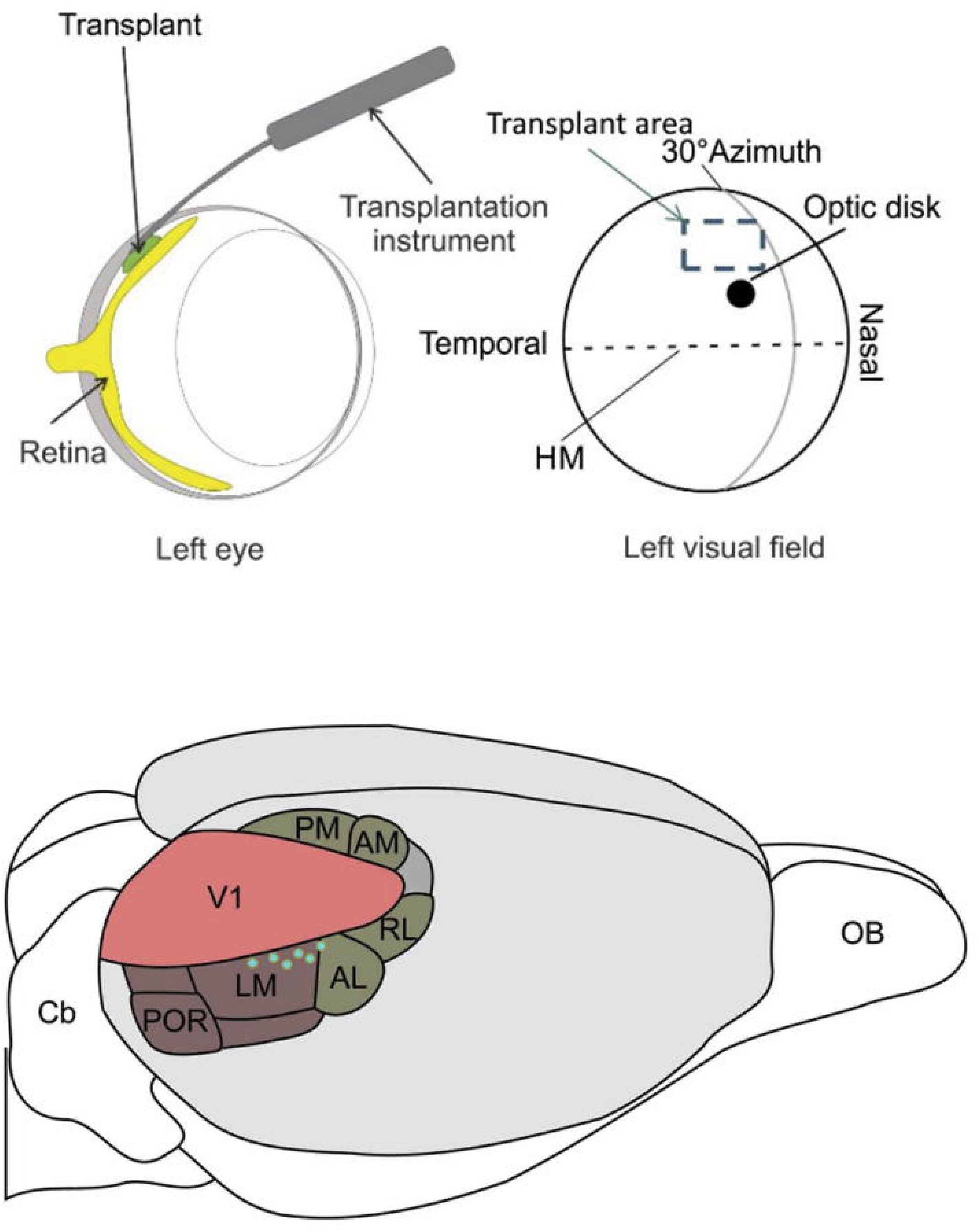
Schematic of experimental setup and approximate place of the retinal sheet transplant. **Top**) retinal sheet transplant derived from fetal transgenic rats is placed in the subretinal space between host-degenerated retina and retinal pigment epithelium using a custom-made surgical tool (Left). The transplant is placed close to the optic disc in the upper temporal area of the visual field (Right). The horizontal meridian (HM) bisects the upper and lower visual field representations. (Adopted from Foik et al 2018). **Bottom**) Approximate recording sites that contained visually selective neurons from transplanted animals. (schematic modified from Scholl et all 2018)

*Optical coherence tomography*. To evaluate transplant placement and development *in vivo*. rats’ eyes were imaged 2-4 weeks after surgery (and then every 2^nd^ month), by spectral domain optical coherence tomography (SC-OCT) using an Envisu R2200 Spectral Domain Ophthalmic Imaging System (Bioptigen, Research Triangle Park, NC). Anesthesia (Seiler et al., 2017) was induced with ketamine/xylazine as above and maintained using isoflurane (1%–2%) mixed with O_2_ through a gas anesthesia mask (Stoelting) if necessary. Pupils were dilated by 1% atropine sulfate ophthalmic solution. Eyes were kept moist between scans with Systane eye drops (Alcon Laboratories). Imaging was accomplished using rectangular scans of a 2.6mm x 2.6mm area at an imaging depth of 1.6 mm. Retinal scans were acquired using at least one of four parameters: 488 B-scans x 488 A-scans x 5 B scan averaging for fundus images; and 800x20x80 for cross-sectional images (units are no. of B scans/no. of A-scans/ B-scan averaging value). Multiple B-scans and fundus images were obtained from different positions to determine graft location, quality of engraftment and state of post-surgery gross retinal architecture. The optic disc was used as a point of reference for locating the transplant and assessing surgical success. Rats exhibiting excess surgical trauma (e.g., choroid damage, optic nerve damage), transplant misplacement (epiretinal, choroid) or corneal opacity/lens cataract were excluded from further analysis.

*Optokinetic Testing (OKT).* After at least 1 hour dark-adaption, the visual acuity of transplanted and non-surgery AMC control rats was measured by recording optomotor responses to a virtual cylinder with alternating black and white vertical stripes at 6 different spatial frequencies (Optomotry, Cerebral Mechanics, Alberta, Canada) at 1, 2, 3, 4 and 5 months post-surgery (MPS), as previously described (McLelland et al., 2018; Seiler et al., 2017). Tests were videotaped and evaluated off-line by two independent observers blinded to the experimental condition. The best visual acuity of the two same-day tests was used for analysis. If there was a discrepancy between the two observers, videos were re-analyzed by a third observer. All testers and video-watchers were blinded to the experimental group of rats. Transplanted rats with poor optokinetic results were deemed unsuitable for recording and were excluded.

*Single-Unit Recording.* Recordings were performed using methods similar to our previous work (Foik et al., 2018). Briefly, rats were anesthetized using 1.5-2% isoflurane in a mixture of N_2_O/O_2_ and a custom designed 3D printed plastic chamber was implanted on the exposed skull using Vetbond™ (3M) and dental acrylics. Animals were immediately transferred to the recording rig. A 4x2 mm craniotomy was made over higher visual areas on the lateral side parallel to skull’s midline. All recordings were performed in the right hemisphere since all retinal transplants were performed on the left eye. After craniotomy and during recordings lighter anesthesia (0.3-0.6%) for the duration of recording was maintained. During this time, ECG and body temperature were monitored. Body temperature was maintained at 37.5 degrees C, using a heating pad underneath the animal (Harvard Apparatus). Single-cell extracellular recordings were made with commercial epoxy-insulated tungsten microelectrodes (5-7 MU, FHC, Bowdoin, ME). To increase chances of finding visually responsive neurons in transplanted and degenerated rats we occasionally used a 32-channel microelectrode array which consisted of 32 single electrodes arranged in 8×4 matrix (MicroProbes, MD, USA). Electrodes were aligned perpendicular to the cortical surface Once the electrode was inserted, the chamber surrounding the craniotomy was filled with 1.5% agar solution in saline and sealed with physiological wax (melting point 45 °C) to reduce brain pulsation. Electrodes were advanced through the cortex by a computer-controlled micro positioner (Motion Controller SMC-100, New-port) fixed to a stereotaxic arm (KOPF Instruments).

At the end of recording sessions, ketamine-xylazine (90 mg/kg Ketamine and 9 mg/kg Xylazine, i.p.) was administered. Animals were then perfused with PFA (4%), and brains and eyes were removed for histology.

*Visual Stimulation.* Data was acquired using a 32-channel Scout recording system. The spike signal was bandpass filtered from 500 Hz to 7 kHz and stored in a computer hard drive at 30 kHz sampling frequency. Spikes were sorted online in Trellis while performing visual stimulation. Visual stimuli were generated in MATLAB (The MathWorks) using Psychophysics Toolbox (Foik et al., 2018) and displayed on a gamma-corrected LCD monitor (48 inches, 120 Hz) at resolution 1920 x 1080 pixels and 52 cd/m 2 mean luminance. For recordings of visually evoked responses, cells were first tested for visual responsiveness with 100 repetitions of a 500 ms bright flash stimulus (105 cd/m 2). Receptive fields for visually responsive cells were located using square-wave drifting gratings, after which optimal orientation/direction and spatial and temporal frequencies were determined using sine wave gratings. Spatial frequencies tested were from 0.001 to 0.5 cycles/°. Temporal frequencies tested were 0.1–10 cycles/s. With these optimal parameters, size tuning was assessed using sizes of 1°-110°, and 100% contrast. With the optimal size, temporal and spatial frequencies, and at high contrast, the orientation selectivity of the cell was tested again using 16 directions stepped by 22.5° increments with full-field size as well as the CRF size. This was followed by testing of contrast.

*Data analysis*. Tuning curves were calculated based on average spike rate. Optimal visual parameters were chosen as the maximum response value.

For tuning width, the orientation responses were fitted to sum of two Gaussian distributions (Alitto & Usrey, 2004; Carandini & Ferster, 2000; Y. J. Liu et al., 2015; Y. J. Liu, Hashemi-Nezhad, & Lyon, 2017) using the following:

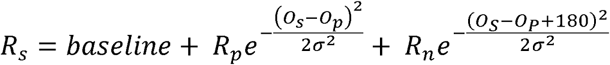

where O_s_ is the stimulus orientation, R_Os_ is the response to different orientations, O_p_ is the preferred orientation, R_p_ and Rn are the responses at the preferred and nonpreferred direction, is the tuning width, and baseline is the offset of the Gaussian distribution. Gaussian fits were estimated without subtracting spontaneous activity, similar to the procedures of (Alitto & Usrey, 2004). The orientation tuning bandwidth of each tuning curve was measured in degrees as the half-width at half-height (HWHH), which equals 1.18×σ based on the equation above.

Size tuning curves were fitted by a difference of Gaussian function (Y. J. Liu, Hashemi-Nezhad, & Lyon, 2011) as follows:

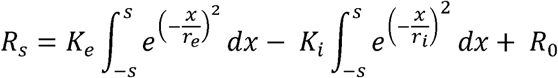

in which R_s_ is the response evoked by different aperture sizes. The free parameters, K_e_ and r_e_, describe the strength and the size of the excitatory space, respectively; K_i_ and R_i_ represent the strength and the size of the inhibitory space, respectively; and R_0_ is the spontaneous activity of the cell.

*Support Vector Machine Classification:* we created a support-vector machine classifier (SVM) object using Matlab ‘s Statistic and Machine Learning Toolbox 12.4 functions (fitcsvm, linear kernel, 10-fold cross-validation). We created a “real” and a null population using same data set with data labels randomly shuffled in each iteration (10 iterations). To reduce classifier bias due to unequal sample sizes of experimental groups, in each iteration, 30 cells from each experimental group were randomly selected to train classifier. Classification results are expressed as balanced accuracy:

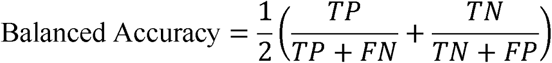

where TP = true positives, FN = false negatives, TN = true negatives, FP = false positives. Statistical significance was assessed using student *t*-test (α = 0.95).

*Statistical analysis*. All statistical comparisons were performed for single-cell populations in three experimental groups: normal, transplant, and degenerated. For each tested parameter, data distributions are represented in two ways: histograms and CDFs. For all tests, results were considered statistically significant at p<0.05. The Kolmogorov–Smirnov test was used for comparisons of distributions and CDFs. A two-tailed Mann–Whitney U test was used for average differences between groups. Mean values given in Results include the SD, and histograms include error bars for the SEM. All offline data analysis and statistics were performed in MATLAB 2022 (MathWorks, Inc.).

### Histology and Immunofluorescence

After euthanasia with injection of anesthetic overdose, rats were perfusion-fixed with cold 4% paraformaldehyde in 0.1 M Na-phosphate buffer. After opening the cornea, eyes were post-fixed overnight at 4°C, then washed. Eye cups were dissected along the dorso-ventral axis, cryoprotected (30% sucrose) and frozen in O.C.T. compound. Serial 10µm cryostat sections were stored at -20°C. Every fifth slide was stained using hematoxylin and eosin (H&E) and imaged on an Olympus BXH10 using an Infinity 3-1U camera. For immunofluorescence a analysis, cryostat sections underwent antigen retrieval at 70 °C with Histo-VT One (Nacalai USA Inc., San Diego, CA), followed by PBS washing, blocking with 20% goat serum, and primary antibodies overnight at 4°C. Slides were incubated for 30-60 min at room temperature in fluorescent or biotinylated secondary antibodies. Primary and secondary antibodies are listed in **Supplemental Table 1**. Fluorescent sections were coverslipped using Vectashield mounting media (Vector Labs, Burlingame, CA) with 5 μg/mL DAPI (4,6-Diamidino-2-phenylindole).

A Leica SP8 confocal microscope (Leica Microsystems) was used for imaging (tiled stacks of 5-8 micron thickness at 20x). Images of confocal stack projections were then extracted using Leica software.

## Results

Transplant morphology (**Figs. 2, S1, S2, S3)**

**Fig. 2.**
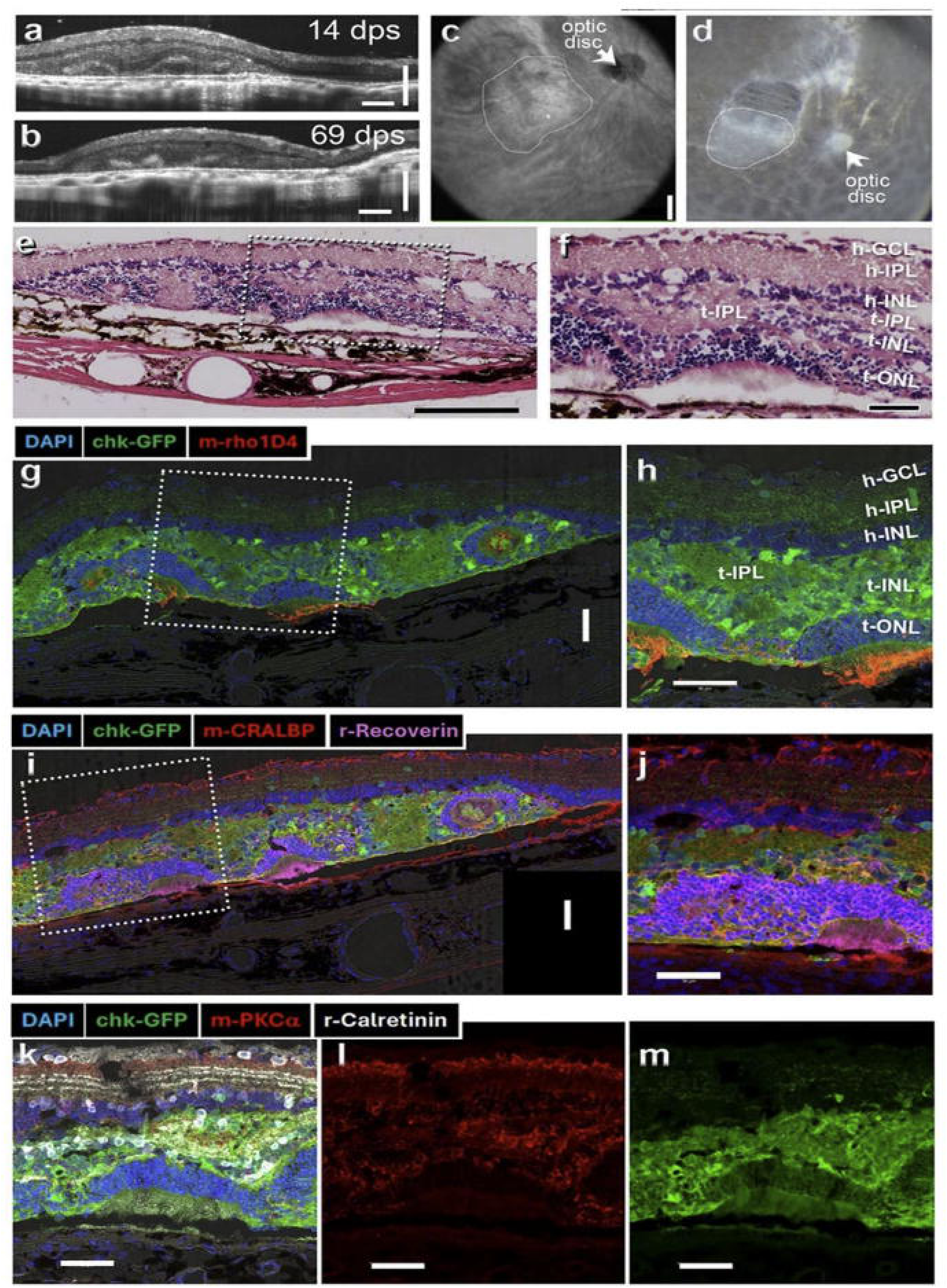
Example of GFP-labeled rat retinal transplant which showed a good response in the visual cortex (135 days post-surgery (dps); rat age 204d). **A), B)** OCT B-scans at 14 and 69 dps Scale bars = 200 µm. **C)** OCT fundus image at 14dps. **D)** transplant in dissected eye cup. **E), F)** H-E staining showing photoreceptor layer in transplant. **G), H)** IHC for rhodopsin and GFP (donor). **I), J)** IHC for recoverin (photoreceptors), CRALBP (glia and RPE) and GFP. **K**), **L**), **M**) IHC for Calretinin, PKCα (rod bipolar cells) and GFP. Scale bars = 200µm (a, b, e); 300 µm (c); 50µm (f-m). Layer labels: h-= host; t-= transplant; GCL = ganglion cell layer; IPL = inner plexiform layer; INL = inner nuclear layer; ONL = outer nuclear layer; OS = outer segment; RPE = retinal pigment epithelium.

**Fig. S1** shows the histology of normal “PA” retina (S1A-C), and an example of degenerated host retina at the age of 204d (**Fig. S1 D-F**), using hematoxylin-eosin (**Fig. S1A,D**), and immunohistochemistry for rhodopsin (rod marker, **Fig. S1B, E**) and recoverin (marker for photoreceptors and cone bipolar cells, **Fig. S1 C,F**). Photoreceptors had mostly degenerated at the time of recording.

SD-OCT confirmed transplant placement into the subretinal space, and distance from optic disc (**Fig. 2A-C; Fig. S2**). Some transplants developed laminated areas, with photoreceptors in contact with host RPE (**Figure S2**, left column), whereas others only developed photoreceptors in rosettes (Figure S2, right column). **Fig. 2** shows an example of a transplant with mixed rosetted and laminated areas that had good cortical responses. In addition, in this transplant, the donor tissue was identified by EGFP expression. Photoreceptors with outer segments could only be found in the transplant (**Fig. 2 G-J**). Transplants were well integrated with the host retina, as shown by staining for rod bipolar cells (PKCα). Figure S3 shows examples of 3 other transplants (without GFP label). Two laminated transplants (transplant 1 with thick outer nuclear layer, Fig. **S3A-D**, and transplant 3 with thin outer nuclear layer, **Fig. S3 E,F**) had good responses, whereas the more disorganized transplant 4 (**Fig. S3 G, H**) showed only weak tuning.

### Retinal sheet transplants significantly increased the number of light-responsive units in V2

We recorded visual responses of the cortical cells in V2 from four treatment groups: healthy controls (“PA” rats, N = 6); line-3 rats with retinal sheet transplants (n =6/10,details in supplementary table 2); line-3rats without transplant (n = 6) and lastly, a group of line-3 rats that underwent a sham transplantation (surgery was performed without transplanting a retinal sheet, n = 3). Since we did not find any selective response in the latter group, no data from this group was included in the rest of analyses in this study. In animals with healthy transplants, 21.03 ± 6.85 per animal (total 112 visual/ 418 no visual) were visually responsive, significantly higher than the occurrence of visually responsive neurons in degenerated rats (p <0.02; **Fig 3A**). Of the 763 well-isolated units recorded in degenerated rats without transplants, only 8.16 ± 3.78% (71), and 2.40 ± 4.94% in rats with sham surgery (2/97). These results show a significant increase in the number of visually responsive units in transplanted animals compared with the degenerated rats (Fig. 3 A-C). Response latencies were measured by briefly flashing the screen on and off 100 times and results are summarized in **Figure 3B**. Neural response latencies were estimated using a Poisson point-process approach. Briefly, spike trains were aligned to stimulus onset, and peri-stimulus time histograms (20 ms bins) were compared against a baseline Poisson distribution of spontaneous activity. Latency was defined as the first two bins in which the spike counts significantly exceeded the 95% confidence bound of baseline firing. On average response latencies of control group (62.9 ± 27.42, n = 434) were significantly shorter than latencies of the transplant and degenerate groups (148.14 ± 113.82, n=86 and 189.15 ± 134.95, n=59 respectively). Also, response latencies of the degenerated group were significantly higher than the transplant group (p<0.01). Moreover, population responses of visually responsive V2 cells transplanted rats showed significant increase in response strength (net response) compared with AMC group (18.80 ± 20.51 and 10.22 ± 6.73 respectively, P<0.01) had much weaker response to screen flashes. Mean net response of the normal controls was measured 52.01 ± 45.32 spikes/second which was very significantly higher that both transplant and AMC groups (**Fig. 3**). Button row panels in **Fig. 3D** provide examples of raster plots from example V2 neurons from control, transplant and degenerated rats, respectively.

**Figure 3.**
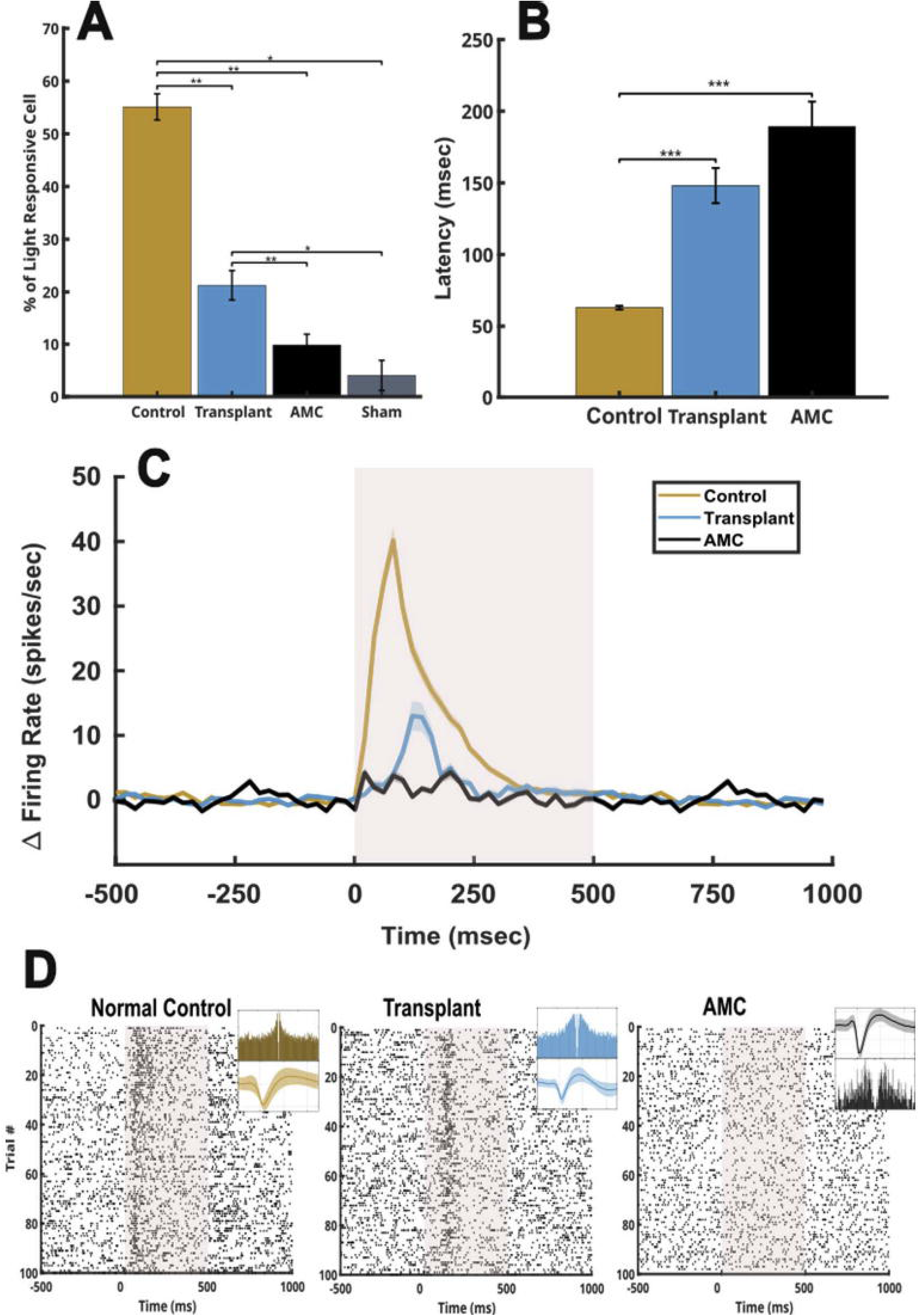
Response latency, population PSTH and proportion of light responsive V2 neurons. **A)** percentages of light responsive cells for each experimental group. **B)** response latency of the three experimental groups after excluding cells from the sham group. **C)** Population PSTH of net firing rates of V2 cells to brief flashes of light across 100 trials. **D)** examples raster plots from the three different treatment groups in response to 100 repetitions of brief flashings of the screen. Boxes inside each represent the spike wave form (top) and autocorrelogram (button) of each isolated unit.

### Size tuning

Size tuning is a key characteristic of visual neurons across the mammals (Y. J. Liu et al., 2017; Van den Bergh et al., 2010) and neural responses to size tuning can be categorized as suppressive which makes up most cells in the primary visual cortex, facilitative and plateau (Y. J. Liu et al., 2011). After finding center of the receptive field and determining preferred temporal and spatial frequencies, we tested cells using a range of different aperture sizes at the center of their receptive field using optimal parameters (1 to 110 visual deg). **Figure 4 A-I** provide three examples of size tuning curves for the treatment groups normal control, transplant and degenerated, respectively. In general, V2 cells from control rats showed significantly higher suppression indices compared with degenerated and transplanted animals (Fig 4K). In examples of control neuron (**Fig. 4A-C**), responses increased with aperture size, peaked at 21°, and decreased for larger apertures, reflecting strong surround suppression.

**Figure 4.**
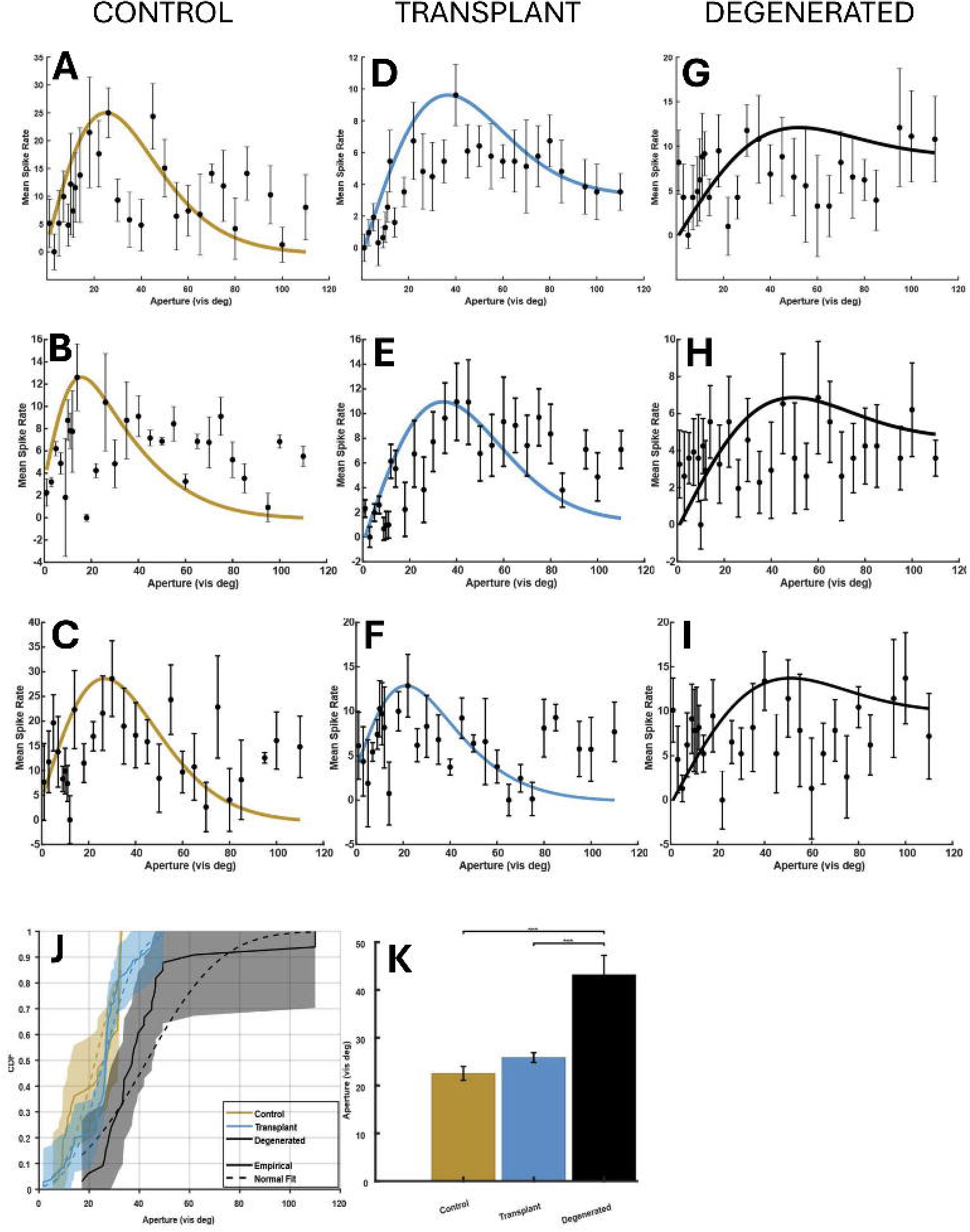
Examples of size tuning curve sand population results. **A-D)** examples of aperture tuning curves of V2 neurons from control rats show relatively strong inhibition compared with neurons from degenerated rats (G-H). **D-F)** examples of aperture tuning curves of V2 neurons from degenerated rats that received retinal transplant in on eye, **G-I)** examples of V2 neurons aperture tuning from degenerated rats. **J)** cumulative distribution functions of size tuning suggest a clear separation of degenerate rats from the two other groups. **K**, mean aperture size for ach treatment group computed from fit (p<0.001).

Examples of transplant neuron (**Fig. 4D-F**) exhibited a similar tuning profile, with clear suppression at larger sizes, although with slightly lower peak firing rates compared to the control examples. By contrast, neurons from the degenerated group (**Fig. 4G-I**) showed a broader tuning curve, characterized by weaker suppression at larger apertures and a higher preferred size, indicating impaired size selectivity. Population-level comparisons are presented in **Fig. 4J-K**. Mean preferred size (**Fig. 4K**) was significantly larger in the degenerated group (44.33° ± 23.39, n = 63) compared with both control (22.19° ± 9.90, n = 47) and transplant (26.53° ± 10.62, n = 106) groups. No significant difference was observed between control and transplant groups. The cumulative distribution of preferred sizes (**Fig. 4J**) further illustrates this pattern, showing a rightward shift in the degenerated group, while control and transplant distributions closely overlapped. Overall, these results demonstrate that V2 cells from control and transplant groups demonstrate similar size tuning characterized by smaller size preference and stronger surround suppression and the degenerated rats differ significantly in those tuning parameters.

### Examples and Population summary of responses to orientation tuning test

Orientation and direction selectivity are prominent characteristics of neurons in primary visual cortex and are key to the processing of more complex visual attributes such as form and motion in higher visual cortex (e.g., (Glickfeld, Histed, & Maunsell, 2013; Glickfeld & Olsen, 2017; Hubel & Wiesel, 1962; Livingstone & Hubel, 1988)). Neurons in V2 exhibit strong extra-classical surround suppression, which enhances contextual modulation by suppressing redundant background input and amplifying salient structure, supporting figure–ground segregation and efficient coding. This mechanism enables V2 to integrate local features into global percepts, bridging low-level vision with mid-level perceptual organization (Hallum & Movshon, 2014; Hashemi-Nezhad & Lyon, 2012; Ziemba, Freeman, Simoncelli, & Movshon, 2018). Given this role, evaluation of extra-classical surround suppression provides a critical measure of the effectiveness of retinal sheet transplants in restoring near-normal vision within the regions of visual space covered by the graft.

After determining preferred spatial and temporal frequencies, we performed a full-field (sizes generally larger than 250°) orientation tuning test. This test was repeated with optimal size, and the center of the classical receptive field (CRF) was determined.

Figure 5A–F illustrates examples of orientation tuning under optimal center-restricted (CRF, solid line) and full-field (FF, dashed line) stimulus conditions across the three experimental groups. In representative control neurons (Fig. 5A, B), enlarging the stimulus to FF size sharpened orientation tuning (HWHH = 17.74° and 18.74° for FF vs. 21.93° and 25.9° for CRF) but simultaneously reduced response strength, consistent with surround suppression. A similar pattern was observed in neurons from the transplant and degenerated groups (Fig. 5C**,D** and **E,F**, respectively), although the magnitude of sharpening and suppression in degenerated examples was notably weaker consistent with prior findings (Foik et al., 2018).

**Figure 5.**
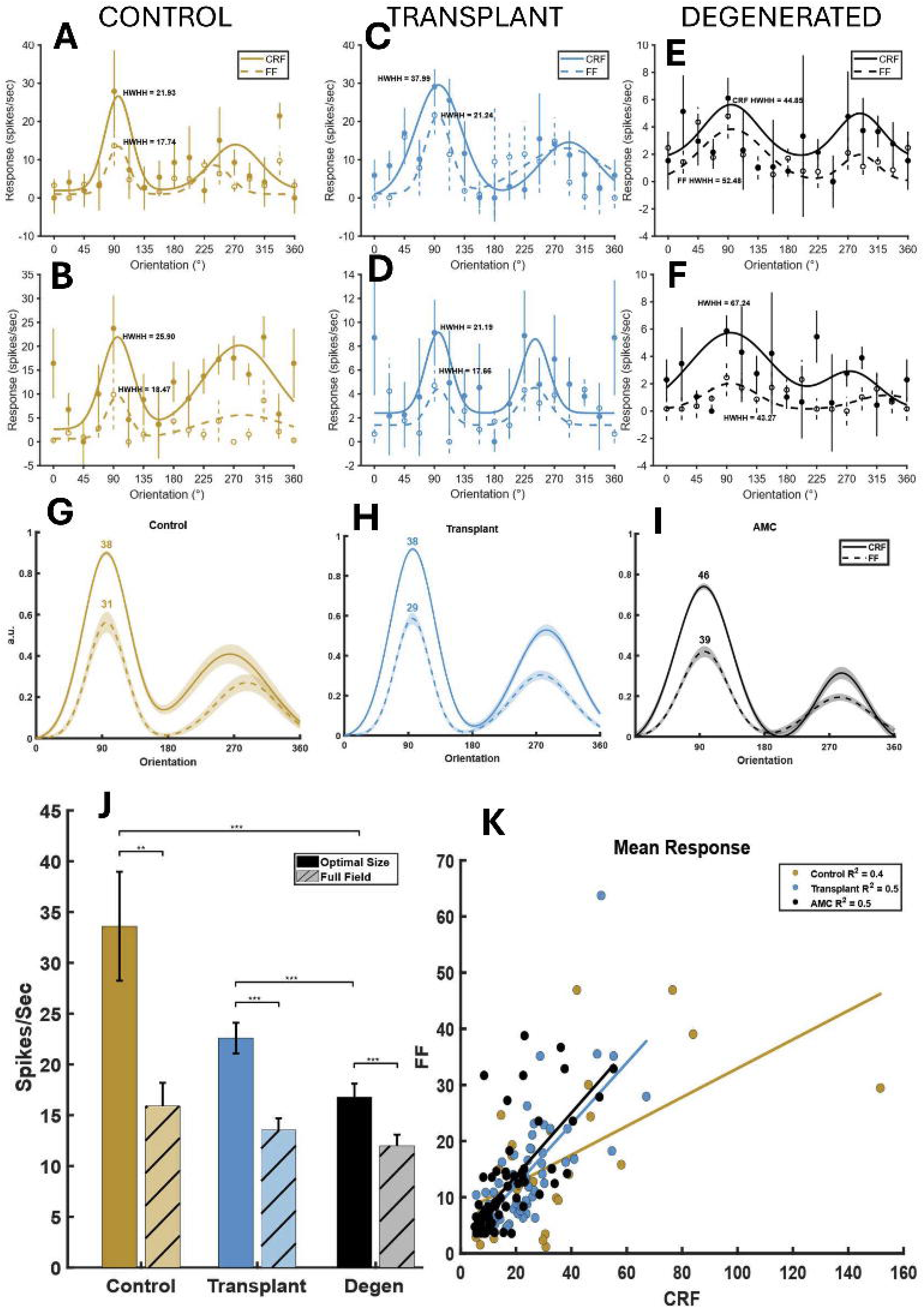
Orientation tuning curves of example cells under full field (FF) and classical receptive field (CRF) stimulus sizes and population response. **A-B, C-D and E-F**, examples of orientation tuning curves in response to stimulus confined to classical receptive field (CRF, solid line) and fullfield stimulus (FF, dashed line) from normal control, transplant and retinal degenerated rats, respectively. **G-I**, The solid line and dashed line represent population of fits from each individual neuron under two size conditions from control, transplant and degenerate rats, respectively. **J**, population response means from all three experimental groups under the two size conditions (p<0.001). **K**, scatter plot of neural responses from all neurons included in this study. All groups show significant linear relationship between responses to CRF and FF stimuli.

Figure 5G–I summarizes the population of calculated tuning curves and HWHH values for each group. Values in these panels represent the HWHH values calculated for the population of cells in each treatment group and stimulus size. While all three groups maintain the same pattern of reduction of firing rate and increase in HWHH as stimulus size gets larger, the overall changes are weaker in the case of degenerated rats.

Fig. 5J illustrates firing rates of V2 neurons in response to their preferred orientation under CRF and FF conditions across Control, Transplant, and Degenerated groups. In Control rats, CRF responses were significantly higher than FF responses (median 27.99 ± 29.39 vs. 13.44 ± 12.53 spikes/sec, Wilcoxon rank-sum, p = 0.0012, z = 3.23). Similarly, in Transplant retinas, CRF responses exceeded FF responses (21.60 ± 12.56 vs. 10.84 ± 9.61 spikes/sec, p < 0.0001, z = 5.27), and in Degenerated (AMC) retinas, CRF responses were greater than FF (12.92 ± 10.87 vs. 8.69 ± 8.93 spikes/sec, p = 0.0007, z = 3.4). Comparisons across groups showed that CRF responses in Control were not significantly different from Transplant (p = 0.0730, z = 1.79) but were significantly higher than Degenerated (p = 0.0004, z = 3.532). FF responses did not differ significantly between Control and Transplant (p = 0.4671, z = 0.73) or between Control and Degenerated (p = 0.1612, z = 1.4), while Transplant FF responses trended higher than Degenerated (p = 0.1044, z = 1.62).

We observed a robust positive correlation between CRF and FF firing rates to preferred orientation across all experimental groups, indicating that surround suppression remains consistent across stimulus sizes (Fig. 5K). These findings suggest that, when visual input is present, V2 neurons preserve both response patterns and surround suppression mechanisms under varying stimulus conditions. These results demonstrate that while transplanted animals exhibit neural activity patterns similar to controls, the retinal degeneration group exhibits a distinct shift that suggests retinal sheet transplantation significantly improved neural responses.

### Population summary of HWHH and Orientation Selectivity Index

Population analyses further revealed group differences in tuning properties. We computed half-width at half-height for responses to the preferred orientation for stimuli confined to the classical receptive field (optimal size) of the cell and very large aperture (full field). As evidenced by HWHH values, tuning curves of all three experimental groups were significantly broadened as stimulus size was reduced from FF to CRF (Fig. 6A). The HWHH tuning curve as an inverse function of stimulus size has been documented in cat (Y. J. Liu et al., 2017), and we are reporting this for the first time in a rodent model. Also, average HWHH of the degenerated group under both stimulus sizes was significantly larger than those of normal control and transplant groups, and no significant difference was observed between control and transplant. This result is similar to our previous findings in V1 of rats with the same treatments (Foik et al., 2018).

**Figure 6.**
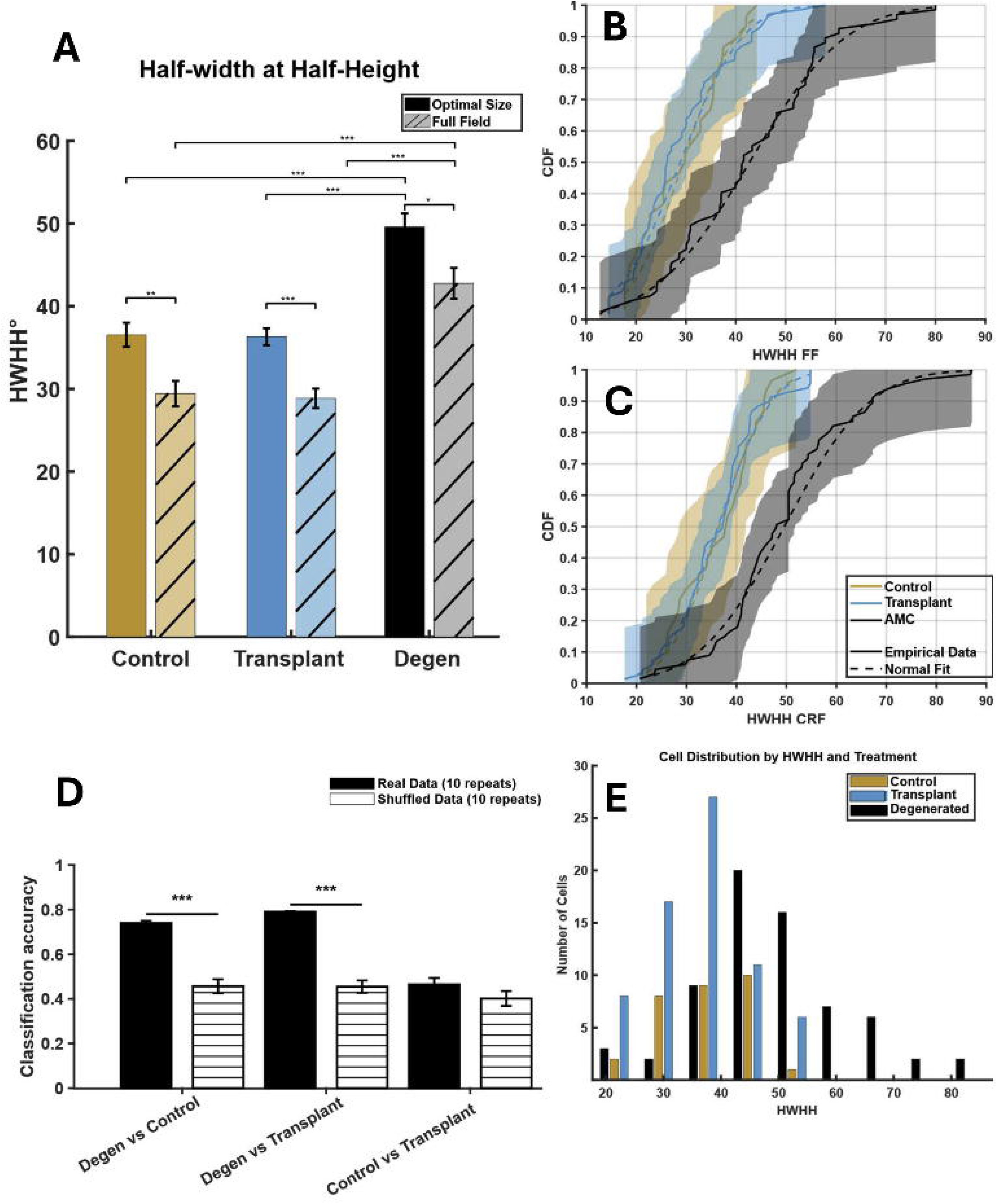
Population Orientation tuning. **A)** HWHH results for the tree treatment groups and under CRF and FF conditions show all groups had significantly sharper tunings under FF and degenerated group had significantly broader tuning under both FF and CRF conditions. (Wilcoxon rank sum test, p<0.05). **B-C)** CDF plots show under both CRF and FF conditions, HWHH values of the degenerated group is clearly separated from the two other. **D)** results of SVM classification using full-filed HWHH and CRF HWHH as input variables. For each iteration, 30 cells were randomly selected for real (solid black bars) and shuffle led (striped bars) data. Shuffled data represent actual chance level for each category. **E)** Distribution of number of cells suggest all treatment groups had relatively normal distribution, but degenerated group contained several cells with HWHH value above 50° in comparison to two other groups. HWHH values taken from CRF condition.

Figure 6A summarizes population half-width at half-height (HWHH) across the three experimental groups. Across all groups, population averages revealed a significant reduction in HWHH under full-field (FF) compared with center-restricted (CRF) stimulation (control: z = 3.06, p = 0.002; transplant: z = 4.62, p ≈ 0; degenerated: z = 2.44, p = 0.014). The degenerated group exhibited significantly broader tuning compared with controls under both CRF (z = –4.80, p ≈ 0) and FF conditions (z = –4.21, p ≈ 0). A similar pattern was observed in the transplant group, which also displayed sharper tuning relative to degenerated animals under both CRF (z = –6.33, p ≈ 0) and FF stimulation (z = –5.59, p ≈ 0). By contrast, average HWHH values did not differ significantly between control and transplant groups under either CRF (z = 0.43, p = 0.67) or FF conditions (z = 0.46, p = 0.64). Consistent with these results, the distribution of HWHH values (Fig. 6E) shows that the degenerated group was clearly shifted toward broader tuning compared with both control and transplant groups.

CDF functions for HWHH in FF and CRF conditions, illustrated in Figure 6B and **6C**, respectively, confirm the aforementioned separation between tuning properties of V2 neurons of degenerated rats and the two other experimental groups.

To further assess group discriminability, we performed pairwise support vector machine (SVM) classification with a linear kernel using HWHH values under the two stimulus sizes as the predictor variables. Classification results are presented in Fig. 6D as classification accuracy. To test real chance level for each treatment pair, we randomly scrambled data labels and created a null distribution (Fig. 6D, striped bars). The classification accuracy for Control vs. Degenerated groups was well above chance (0.74 ± 0.031 vs. null: 0.457 ± 0.099, p ≈ 0), indicating robust separability of these populations. Similarly, the SVM classifier was able to separate the tuning widths of the Degenerated and Transplant groups with remarkable accuracy. Here, classification accuracy reached 0.790 ± 0.008, far above null expectations (0.455 ± 0.087, p ≈ 0), reflecting very significant divergence between the two populations. By contrast, Control vs. Transplant classification did not exceed chance performance (0.465 ± 0.091 vs. 0.402 ± 0.106, p = 0.4), suggesting that population-level activity patterns in transplanted animals closely resemble those of controls. Consistent with these findings, the distribution of HWHH values showed substantial overlap between Control and Transplant groups, but clear separation of Degenerated animals from both. Collectively, these results demonstrate that surround suppression sharpens orientation tuning in V2 neurons and that this mechanism is largely preserved in transplant recipients but substantially degraded in degenerated animals. We concluded that: 1) rat V2 neurons exhibit surround suppression profile similar to what was reported in cats (Hashemi-Nezhad & Lyon, 2012) and monkeys (Hallum & Movshon, 2014), 2) surround suppression is present across all three treatment groups and 3) retinal degeneration results in much broader tuning curves compared to normal control and that retinal transplant significantly improves tuning widths.

## Discussion

In this study, we show that retinal sheet transplants can lead to nearly normal quality visual responses at the cortical level in animal models with severe retinal degeneration. Transplants not only helped to improve sight well beyond the capabilities of non-transplanted animals but also preserved underlying neural connectivity. This method shows promise for treating advanced stages of retinal diseases, such as macular degeneration and retinitis pigmentosa, where most photoreceptors are gone, and for which no treatments currently exist.

The fetal retinal sheet transplant is a well-organized structure of fetal progenitor cells, which differentiate into fully functional photoreceptors that integrate with bipolar and amacrine cells of the host retina. The reliability of retinal sheet transplants in the manner used here has been verified now in several previous animal studies (Aramant & Seiler, 2002; Seiler & Aramant, 1998; Seiler et al., 2017; Seiler et al., 2005; Yang et al., 2010) and even in humans (Radtke et al., 2008). In this study, we verified that transplants were well integrated into the host retina and produced new photoreceptors and rod bipolar cells. While we did not directly confirm synaptic connectivity in this study, elsewhere this has been confirmed repeatedly for this same type of transplant in this same rat model (Seiler et al., 2010; Seiler, Thomas, Chen, Wu, et al., 2008). Here we found that, within months of receiving the retinal sheet transplant, V1 neurons in line-3-degenerated rats responded similarly to normal rats, revealing a high degree of selectivity to orientation, direction, spatial and temporal frequency, and contrast. In degenerated rats that did not receive a transplant, significantly fewer neurons were visually responsive and exhibited poor selectivity to all stimulus categories. This is consistent with other studies using non-transplanted line-3 rats where visual stimuli did not elicit a response in most neurons of V1 (Chen et al., 2016) and the superior colliculus (Seiler, Thomas, Chen, Arai, et al., 2008; Seiler, Thomas, Chen, Wu, et al., 2008; Thomas, Seiler, Sadda, & Aramant, 2004; Yang et al., 2010). To be clear, these studies, as well as ours, do report some visually responsive neurons in degenerated rats. This is likely due to some remaining cones, which have been detected as late as P90 in line-3 rats (Ray et al., 2010). The number of visually responsive cells in transplanted rats was higher than in degenerated rats, but lower than in normal rats. Nevertheless, responsive neurons were highly selective, indicating a good degree of normal visual function.

Furthermore, the location of nonresponsive neurons correlated with retinotopic regions of visual cortex that lay at the fringe or outside of the region covered by the retinal sheet transplant. Similar results in response to flashes of light were reported for neurons in the superior colliculus (Seiler, Thomas, Chen, Arai, et al., 2008; Thomas et al., 2004; Yang et al., 2010). This indicates that the generation of visual responses is due to the transplant itself; and, furthermore, that more complete recovery of V2 function would be possible through transplants of sheets covering more retinal area. There is a possibility that transplants led to neuroprotection of the remaining host photoreceptors, which also contributed to improved cortical responses. However, studies using line-3 rats have found that remaining host photoreceptors were similar in number regardless of proximity to the fetal tissue graft, suggesting that there is not a neuroprotective effect (Seiler & Aramant, 1998; Seiler et al., 2017).

Another of our findings was that response latency was longer for transplanted rat V2 neurons compared with normal rats. A similar increase in latency from normal rats was found using line-3 rats while recording from the superior colliculus (Yang et al., 2010), which shows that the lag is not exclusive to cortical neurons. Possible reasons for the increased latency include that the summing of inputs from photoreceptors in the transplanted retina through to ganglion cells of the original retina has either longer to travel due to the added layer of retina or longer to summate due to a weaker level of photoreceptor output. Nevertheless, the delay does not seem to perturb selectivity of V2 neurons in transplanted rats as we observed.

In our previous study (Foik et al., 2018) using viral tracing with monosynaptic rabies virus we demonstrated that feed-forward inputs from visual thalamus, connectivity within V1, and feedback from higher visual areas were present in degenerated rats, but overall, to a lesser extent than transplanted and normal rats. This is important because it reveals that underlying visual cortical circuitry is available for retinal sheet transplants to engage. Furthermore, our findings show that transplantation restores the circuitry to a level comparable with normal rats. In our current study, we have proved that cortical circuitry, especially at the higher visual cortical level, is functional even after end stages of retinal degeneration results in near complete loss of visual inputs and can process visual inputs at qualities comparable to healthy controls.

In conclusion, we showed that vision loss in an animal model of severe retinal degeneration can be remedied, and normal circuitry and function can be restored in higher visual cortex. Fetal retinal sheet transplants can be successfully incorporated into the host tissue to deliver the signals necessary for highly selective visual responses in cortex. These results show the potential this approach may have to restore vision in people suffering from late-stage macular degeneration or retinitis pigmentosa.

## Supporting information

sad

## Acknowledgements

Authors declare no conflicts of interest.

This study was supported by the National Eye Institute grants R01EY032948 and R01EY031834. This work was also made possible, in part, through access to the Optical Biology Core Facility of the Developmental Biology Center, a shared resource supported by the Cancer Center Support Grant (CA-62203) and Center for Complex Biological Systems Support Grant (GM-076516) at the University of California, Irvine. The authors acknowledge support to the Gavin Herbert Eye Institute at the University of California, Irvine from an unrestricted grant from Research to Prevent Blindness and from NIH grant P30 EY034070 to the Department of Ophthalmology & Visual Sciences at UC Irvine.

The authors wish to thank Dr. Andrzej Foik for his contribution to early experimental design and data analysis and the current and previous staff members: Helios Nguyen, Robert Sims, Akashi Suon, Meredith Liu; and the students Tammy Xie, Ceci Zhang, Ani Khachigian, Topper Quimpo, Maihan Phan, Quinn Tran, Natalie Merida Alvarez, Hao Tran, Kelly Tran, Dominic Rojas, Kylie Tran, Sahar Ahmed, Viviana Moreno, Irene Jin, Smeet Shah, Alex Linn, Nick Tang, Jack Wang, Merlin Marianandan, Umer Shahab and many other students for technical assistance.

## Author contributions

AA: performed experiments, performed analysis, wrote the paper, approved final version of manuscript

BL: performed experiments, performed analysis, approved final version of manuscript

MJS: provided funding, performed experiments, performed analysis, wrote the paper, approved final version of manuscript

DCL: provided funding, designed experiments, performed analysis, wrote the paper, approved final version of manuscript

## References

Alitto, H. J., & Usrey, W. M. (2004). Influence of contrast on orientation and temporal frequency tuning in ferret primary visual cortex. J Neurophysiol, 91(6), 2797–2808. doi:10.1152/jn.00943.2003

Aramant, R. B., & Seiler, M. J. (2002). Retinal transplantation--advantages of intact fetal sheets. Prog Retin Eye Res, 21(1), 57–73. doi:10.1016/s1350-9462(01)00020-9

Berson, E. L., Rosner, B., Sandberg, M. A., Weigel-DiFranco, C., Moser, A., Brockhurst, R. J., . . . Schaefer, E. J. (2004). Clinical trial of docosahexaenoic acid in patients with retinitis pigmentosa receiving vitamin A treatment. Arch Ophthalmol, 122(9), 1297–1305. doi:10.1001/archopht.122.9.1297

Bertolotti, E., Neri, A., Camparini, M., Macaluso, C., & Marigo, V. (2014). Stem cells as source for retinal pigment epithelium transplantation. Prog Retin Eye Res, 42, 130–144. doi:10.1016/j.preteyeres.2014.06.002

Carandini, M., & Ferster, D. (2000). Membrane potential and firing rate in cat primary visual cortex. J Neurosci, 20(1), 470–484. doi:10.1523/JNEUROSCI.20-01-00470.2000

Cavanaugh, J. R., Bair, W., & Movshon, J. A. (2002). Selectivity and spatial distribution of signals from the receptive field surround in macaque V1 neurons. J Neurophysiol, 88(5), 2547–2556. doi:10.1152/jn.00693.2001

Chen, K., Wang, Y., Liang, X., Zhang, Y., Ng, T. K., & Chan, L. L. (2016). Electrophysiology Alterations in Primary Visual Cortex Neurons of Retinal Degeneration (S334ter-line-3) Rats. Sci Rep, 6, 26793. doi:10.1038/srep26793

Coffey, P. J., Girman, S., Wang, S. M., Hetherington, L., Keegan, D. J., Adamson, P., . . . Lund, R. D. (2002). Long-term preservation of cortically dependent visual function in RCS rats by transplantation. Nat Neurosci, 5(1), 53–56. doi:10.1038/nn782

Coogan, T. A., & Burkhalter, A. (1993). Hierarchical organization of areas in rat visual cortex. J Neurosci, 13(9), 3749–3772. doi:10.1523/JNEUROSCI.13-09-03749.1993

Fleckenstein, M., Keenan, T. D. L., Guymer, R. H., Chakravarthy, U., Schmitz-Valckenberg, S., Klaver, C. C., . . . Chew, E. Y. (2021). Age-related macular degeneration. Nat Rev Dis Primers, 7(1), 31. doi:10.1038/s41572-021-00265-2

Foik, A. T., Lean, G. A., Scholl, L. R., McLelland, B. T., Mathur, A., Aramant, R. B., . . . Lyon, D. C. (2018). Detailed Visual Cortical Responses Generated by Retinal Sheet Transplants in Rats with Severe Retinal Degeneration. J Neurosci, 38(50), 10709–10724. doi:10.1523/JNEUROSCI.1279-18.2018

Gias, C., Vugler, A., Lawrence, J., Carr, A. J., Chen, L. L., Ahmado, A., . . . Coffey, P. J. (2011). Degeneration of cortical function in the Royal College of Surgeons rat. Vision Res, 51(20), 2176–2185. doi:10.1016/j.visres.2011.08.012

Girman, S. V., Wang, S., & Lund, R. D. (2003). Cortical visual functions can be preserved by subretinal RPE cell grafting in RCS rats. Vision Res, 43(17), 1817–1827. doi:10.1016/s0042-6989(03)00276-1

Glickfeld, L. L., Histed, M. H., & Maunsell, J. H. (2013). Mouse primary visual cortex is used to detect both orientation and contrast changes. J Neurosci, 33(50), 19416–19422. doi:10.1523/JNEUROSCI.3560-13.2013

Glickfeld, L. L., & Olsen, S. R. (2017). Higher-Order Areas of the Mouse Visual Cortex. Annu Rev Vis Sci, 3, 251–273. doi:10.1146/annurev-vision-102016-061331

Glickfeld, L. L., Reid, R. C., & Andermann, M. L. (2014). A mouse model of higher visual cortical function. Curr Opin Neurobiol, 24(1), 28–33. doi:10.1016/j.conb.2013.08.009

Hallum, L. E., & Movshon, J. A. (2014). Surround suppression supports second-order feature encoding by macaque V1 and V2 neurons. Vision Res, 104, 24–35. doi:10.1016/j.visres.2014.10.004

Hashemi-Nezhad, M., & Lyon, D. C. (2012). Orientation tuning of the suppressive extraclassical surround depends on intrinsic organization of V1. Cereb Cortex, 22(2), 308–326. doi:10.1093/cercor/bhr105

Hombrebueno, J. R., Tsai, M. M., Kim, H. L., De Juan, J., Grzywacz, N. M., & Lee, E. J. (2010). Morphological changes of short-wavelength cones in the developing S334ter-3 transgenic rat. Brain Res, 1321, 60–66. doi:10.1016/j.brainres.2010.01.051

Hubel, D. H., & Wiesel, T. N. (1962). Receptive fields, binocular interaction and functional architecture in the cat’s visual cortex. J Physiol, 160(1), 106–154. doi:10.1113/jphysiol.1962.sp006837

Juavinett, A. L., & Callaway, E. M. (2015). Pattern and Component Motion Responses in Mouse Visual Cortical Areas. Curr Biol, 25(13), 1759–1764. doi:10.1016/j.cub.2015.05.028

Kauper, K., McGovern, C., Sherman, S., Heatherton, P., Rapoza, R., Stabila, P., . . . Tao, W. (2012). Two-year intraocular delivery of ciliary neurotrophic factor by encapsulated cell technology implants in patients with chronic retinal degenerative diseases. Invest Ophthalmol Vis Sci, 53(12), 7484–7491. doi:10.1167/iovs.12-9970

Kobatake, E., & Tanaka, K. (1994). Neuronal selectivities to complex object features in the ventral visual pathway of the macaque cerebral cortex. J Neurophysiol, 71(3), 856–867. doi:10.1152/jn.1994.71.3.856

LaVail, M. M., Nishikawa, S., Steinberg, R. H., Naash, M. I., Duncan, J. L., Trautmann, N., . . . Flannery, J. G. (2018). Phenotypic characterization of P23H and S334ter rhodopsin transgenic rat models of inherited retinal degeneration. Exp Eye Res, 167, 56–90. doi:10.1016/j.exer.2017.10.023

Lipinski, D. M., Thake, M., & MacLaren, R. E. (2013). Clinical applications of retinal gene therapy. Prog Retin Eye Res, 32, 22–47. doi:10.1016/j.preteyeres.2012.09.001

Liu, M. M., Tuo, J., & Chan, C. C. (2011). Gene therapy for ocular diseases. Br J Ophthalmol, 95(5), 604–612. doi:10.1136/bjo.2009.174912

Liu, Y. J., Hashemi-Nezhad, M., & Lyon, D. C. (2011). Dynamics of extraclassical surround modulation in three types of V1 neurons. J Neurophysiol, 105(3), 1306–1317. doi:10.1152/jn.00692.2010

Liu, Y. J., Hashemi-Nezhad, M., & Lyon, D. C. (2015). Contrast invariance of orientation tuning in cat primary visual cortex neurons depends on stimulus size. J Physiol, 593(19), 4485–4498. doi:10.1113/JP271180

Liu, Y. J., Hashemi-Nezhad, M., & Lyon, D. C. (2017). Differences in orientation tuning between pinwheel and domain neurons in primary visual cortex depend on contrast and size. Neurophotonics, 4(3), 031209. doi:10.1117/1.NPh.4.3.031209

Livingstone, M., & Hubel, D. (1988). Segregation of form, color, movement, and depth: anatomy, physiology, and perception. Science, 240(4853), 740–749. doi:10.1126/science.3283936

Marshel, J. H., Garrett, M. E., Nauhaus, I., & Callaway, E. M. (2011). Functional specialization of seven mouse visual cortical areas. Neuron, 72(6), 1040–1054. doi:10.1016/j.neuron.2011.12.004

Martinez-Navarrete, G., Seiler, M. J., Aramant, R. B., Fernandez-Sanchez, L., Pinilla, I., & Cuenca, N. (2011). Retinal degeneration in two lines of transgenic S334ter rats. Exp Eye Res, 92(3), 227–237. doi:10.1016/j.exer.2010.12.001

McLelland, B. T., Lin, B., Mathur, A., Aramant, R. B., Thomas, B. B., Nistor, G., . . . Seiler, M. J. (2018). Transplanted hESC-Derived Retina Organoid Sheets Differentiate, Integrate, and Improve Visual Function in Retinal Degenerate Rats. Invest Ophthalmol Vis Sci, 59(6), 2586–2603. doi:10.1167/iovs.17-23646

Radtke, N. D., Aramant, R. B., Petry, H. M., Green, P. T., Pidwell, D. J., & Seiler, M. J. (2008). Vision improvement in retinal degeneration patients by implantation of retina together with retinal pigment epithelium. Am J Ophthalmol, 146(2), 172–182. doi:10.1016/j.ajo.2008.04.009

Ray, A., Sun, G. J., Chan, L., Grzywacz, N. M., Weiland, J., & Lee, E. J. (2010). Morphological alterations in retinal neurons in the S334ter-line3 transgenic rat. Cell Tissue Res, 339(3), 481–491. doi:10.1007/s00441-009-0916-5

Sagdullaev, B. T., Aramant, R. B., Seiler, M. J., Woch, G., & McCall, M. A. (2003). Retinal transplantation-induced recovery of retinotectal visual function in a rodent model of retinitis pigmentosa. Invest Ophthalmol Vis Sci, 44(4), 1686–1695. doi:10.1167/iovs.02-0615

Schwartz, S. D., Regillo, C. D., Lam, B. L., Eliott, D., Rosenfeld, P. J., Gregori, N. Z., . . . Lanza, R. (2015). Human embryonic stem cell-derived retinal pigment epithelium in patients with age-related macular degeneration and Stargardt’s macular dystrophy: follow-up of two open-label phase 1/2 studies. Lancet, 385(9967), 509–516. doi:10.1016/S0140-6736(14)61376-3

Seiler, M. J., & Aramant, R. B. (1998). Intact sheets of fetal retina transplanted to restore damaged rat retinas. Invest Ophthalmol Vis Sci, 39(11), 2121–2131.

Seiler, M. J., Aramant, R. B., & Ball, S. L. (1999). Photoreceptor function of retinal transplants implicated by light-dark shift of S-antigen and rod transducin. Vision Res, 39(15), 2589–2596. doi:10.1016/s0042-6989(98)00326-5

Seiler, M. J., Aramant, R. B., Jones, M. K., Ferguson, D. L., Bryda, E. C., & Keirstead, H. S. (2014). A new immunodeficient pigmented retinal degenerate rat strain to study transplantation of human cells without immunosuppression. Graefes Arch Clin Exp Ophthalmol, 252(7), 1079–1092. doi:10.1007/s00417-014-2638-y

Seiler, M. J., Aramant, R. B., Thomas, B. B., Peng, Q., Sadda, S. R., & Keirstead, H. S. (2010). Visual restoration and transplant connectivity in degenerate rats implanted with retinal progenitor sheets. Eur J Neurosci, 31(3), 508–520. doi:10.1111/j.1460-9568.2010.07085.x

Seiler, M. J., Lin, R. E., McLelland, B. T., Mathur, A., Lin, B., Sigman, J., . . . Thomas, B. B. (2017). Vision Recovery and Connectivity by Fetal Retinal Sheet Transplantation in an Immunodeficient Retinal Degenerate Rat Model. Invest Ophthalmol Vis Sci, 58(1), 614–630. doi:10.1167/iovs.15-19028

Seiler, M. J., Sagdullaev, B. T., Woch, G., Thomas, B. B., & Aramant, R. B. (2005). Transsynaptic virus tracing from host brain to subretinal transplants. Eur J Neurosci, 21(1), 161–172. doi:10.1111/j.1460-9568.2004.03851.x

Seiler, M. J., Thomas, B. B., Chen, Z., Arai, S., Chadalavada, S., Mahoney, M. J., . . . Aramant, R. B. (2008). BDNF-treated retinal progenitor sheets transplanted to degenerate rats: improved restoration of visual function. Exp Eye Res, 86(1), 92–104. doi:10.1016/j.exer.2007.09.012

Seiler, M. J., Thomas, B. B., Chen, Z., Wu, R., Sadda, S. R., & Aramant, R. B. (2008). Retinal transplants restore visual responses: trans-synaptic tracing from visually responsive sites labels transplant neurons. Eur J Neurosci, 28(1), 208–220. doi:10.1111/j.1460-9568.2008.06279.x

Self, M. W., Lorteije, J. A., Vangeneugden, J., van Beest, E. H., Grigore, M. E., Levelt, C. N., . . . Roelfsema, P. R. (2014). Orientation-tuned surround suppression in mouse visual cortex. J Neurosci, 34(28), 9290–9304. doi:10.1523/JNEUROSCI.5051-13.2014

Thomas, B. B., Aramant, R. B., Sadda, S. R., & Seiler, M. J. (2005). Light response differences in the superior colliculus of albino and pigmented rats. Neurosci Lett, 385(2), 143–147. doi:10.1016/j.neulet.2005.05.034

Thomas, B. B., Seiler, M. J., Sadda, S. R., & Aramant, R. B. (2004). Superior colliculus responses to light - preserved by transplantation in a slow degeneration rat model. Exp Eye Res, 79(1), 29–39. doi:10.1016/j.exer.2004.02.016

Tsai, Y., Lu, B., Bakondi, B., Girman, S., Sahabian, A., Sareen, D., . . . Wang, S. (2015). Human iPSC-Derived Neural Progenitors Preserve Vision in an AMD-Like Model. Stem Cells, 33(8), 2537–2549. doi:10.1002/stem.2032

Van den Bergh, G., Zhang, B., Arckens, L., & Chino, Y. M. (2010). Receptive-field properties of V1 and V2 neurons in mice and macaque monkeys. J Comp Neurol, 518(11), 2051–2070. doi:10.1002/cne.22321

Vinje, W. E., & Gallant, J. L. (2000). Sparse coding and decorrelation in primary visual cortex during natural vision. Science, 287(5456), 1273–1276. doi:10.1126/science.287.5456.1273

Woch, G., Aramant, R. B., Seiler, M. J., Sagdullaev, B. T., & McCall, M. A. (2001). Retinal transplants restore visually evoked responses in rats with photoreceptor degeneration. Invest Ophthalmol Vis Sci, 42(7), 1669–1676.

Yang, P. B., Seiler, M. J., Aramant, R. B., Yan, F., Mahoney, M. J., Kitzes, L. M., & Keirstead, H. S. (2010). Trophic factors GDNF and BDNF improve function of retinal sheet transplants. Exp Eye Res, 91(5), 727–738. doi:10.1016/j.exer.2010.08.022

Zhu, C. L., Ji, Y., Lee, E. J., & Grzywacz, N. M. (2013). Spatiotemporal pattern of rod degeneration in the S334ter-line-3 rat model of retinitis pigmentosa. Cell Tissue Res, 351(1), 29–40. doi:10.1007/s00441-012-1522-5

Ziemba, C. M., Freeman, J., Simoncelli, E. P., & Movshon, J. A. (2018). Contextual modulation of sensitivity to naturalistic image structure in macaque V2. J Neurophysiol, 120(2), 409–420. doi:10.1152/jn.00900.2017

