## Supplementary material for "Restoration of Higher-Order Cortical Processing in Rat V2 by Retinal Sheet Transplants in Degenerated Rats": sad

**SUPPLEMENTARY FIGURES**

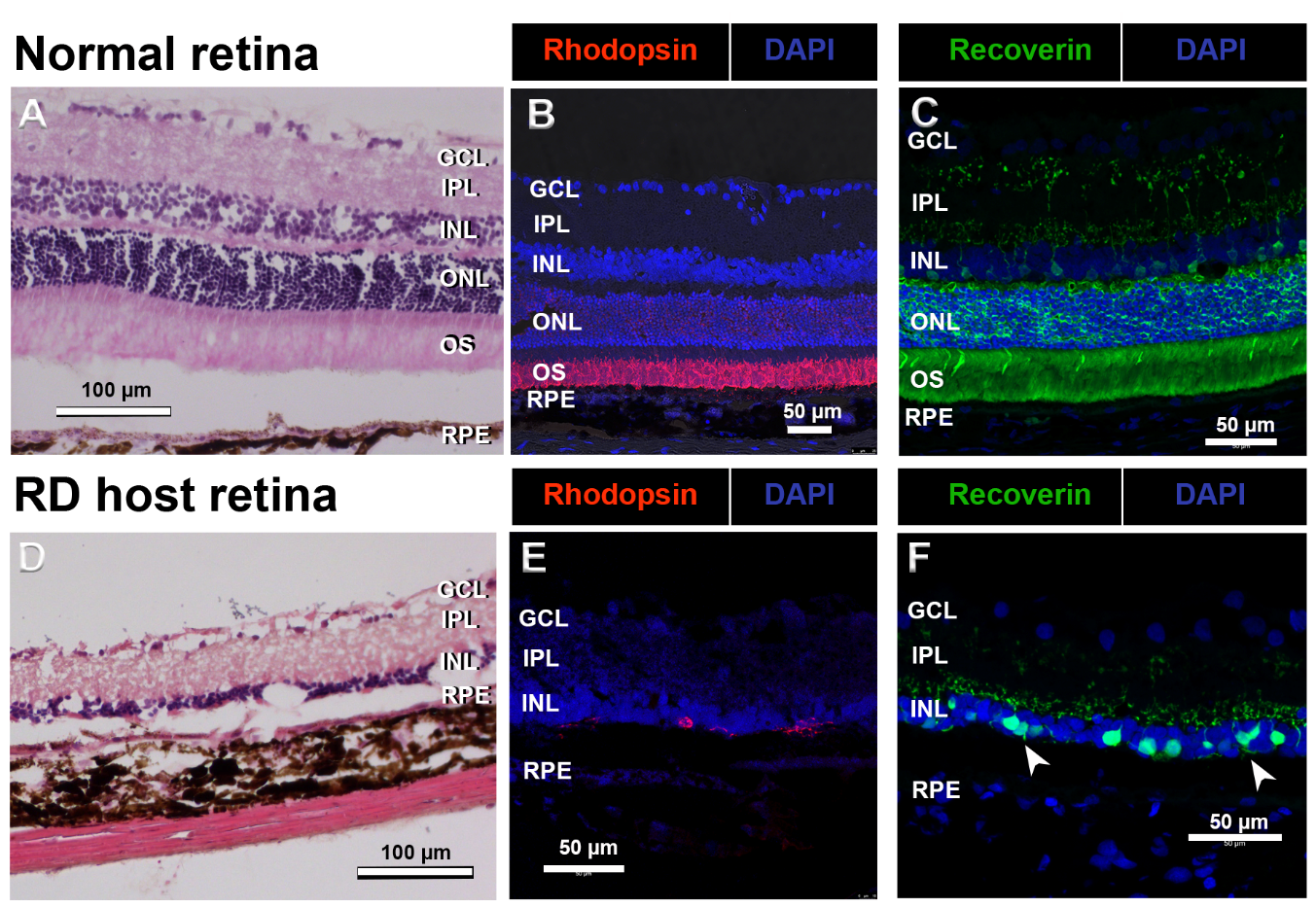

**Supplemental Figure S1**. Comparison of normal rat retina **(A-C)** (rat age 363d (A), 156d (B,C)) with retinal degenerate (RD) host retina (Transplant host in Figure 2, age 204d). **A) D)** Hematoxylin-Eosin staining. Note the difference in retinal thickness between normal and RD retina. **B) E)** Rhodopsin staining of rods. There are very few abnormally looking cells in RD retina. **C) F)** Recoverin (marker for photoreceptors and cone bipolar cells. Most of the recoverin + cells in the RD retina are cone bipolar cells, not photoreceptors. Arrowheads point to degenerating photoreceptors.

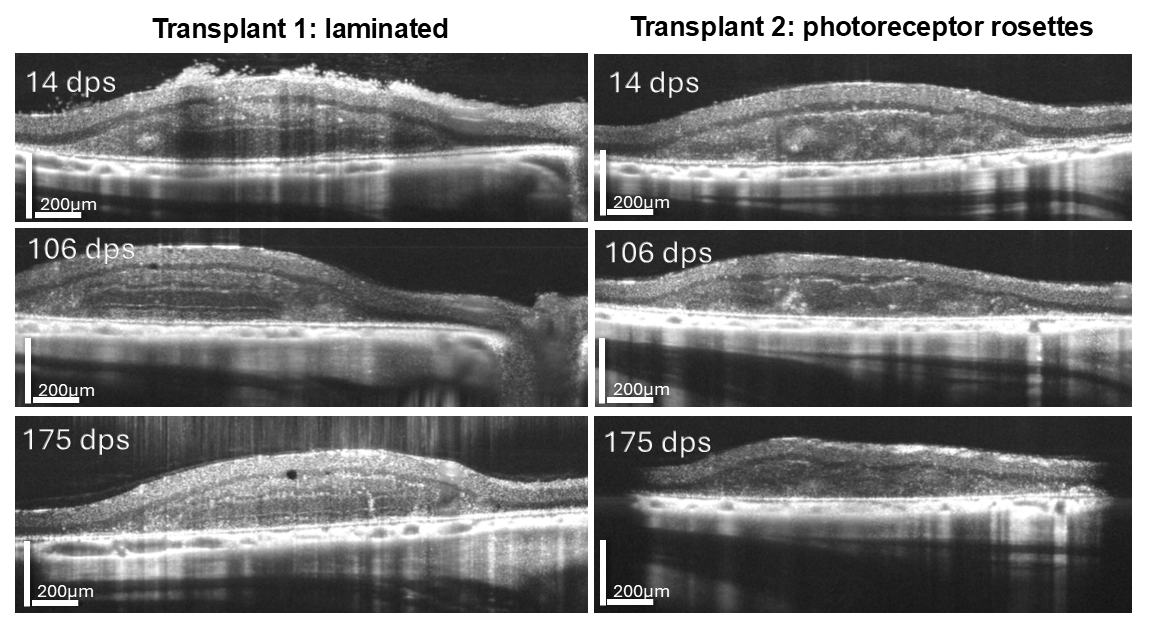

**Supplemental Figure S2. Live imaging using optical coherence tomography.** Comparison of laminated transplant 1 (left column) with rosetted transplant 2 (right column), at 14, 106 and 175 days post-surgery (dps). Transplant 1 had relatively good cortical responses whereas the responses in transplant 2 were weak. Scale = 200 µm.

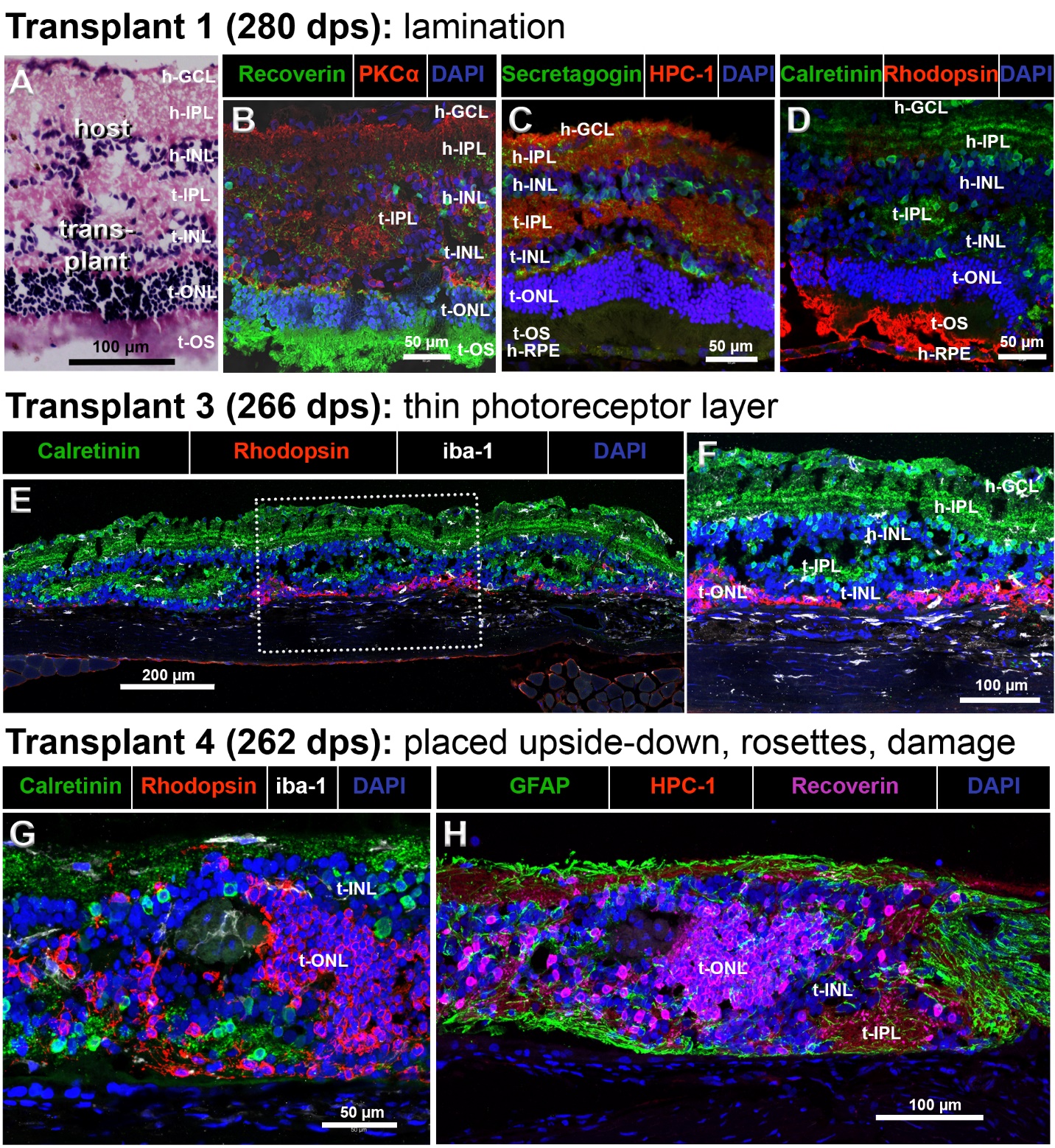

**Supplemental Figure S3. Histology and immunohistochemistry of various transplants.** **A)-D). Transplant 1**, 280dps, rat age 358d. Relatively good cortical responses. This transplant is laminated, with clear outer nuclear layer and photoreceptor outer segments towards host RPE. **A)** H-E staining. The transplant has developed all retinal layers. **B)** Recoverin (green, marker for photoreceptors and cone bipolar cells) in combination with PKCα (red, marker of rod bipolar cells). **C)** Secretagogin (green, marker for cone bipolar cells) in combination with HPC-1/syntaxin (red; label of synaptic layers). **E) F) Transplant 3**, 266dps, rat age 317d. Good cortical responses. **E)** Calretinin (green, ganglion and amacrine cells) in combination with rhodopsin (red, rods), and iba-1 (white, microglia). Very thin photoreceptor layer (red). **F)** Enlargement of box in E). **G,H) Transplant 4**, 262 dps, rat age 312d. Weak tuning responses in cortex. This transplant was placed upside-down into a retinal lesion site (exposed to vitreous) and shows disorganization (rosettes). **G)** Calretinin/rhodopsin/iba-1 (as in E). Note iba1+ cell inside photoreceptor rosette. **H)** Combination of GFAP (green, reactive glia), HPC-1/syntaxin (red, synaptic layers) and recoverin (magenta, photoreceptors). Scale = 50µm (B,C,D,G); 100µm (A,F,H); 200µm (E). Layer labels: h- = host; t- = transplant; GCL = ganglion cell layer; IPL = inner plexiform layer; INL = inner nuclear layer; ONL = outer nuclear layer; OS = outer segment; RPE = retinal pigment epithelium.

| **S1.A. Primary antibodies (IHC)** | | | | | | |
| --- | --- | --- | --- | --- | --- | --- |
| **Antigen** | **species** | **specific for** | **dilution** | **Supplier** | **Catalogue #** | **RRID** |
| Calretinin (SP13) | rabbit | Ganglion and amacrine cells | 1:200 | Thermofisher / Invitrogen | MA5-14540 | AB_10985167 |
| GFAP | goat | Reactive glia | 1:500 | Millipore Sigma | SAb2500462 | AB_10603437 |
| CRALBP | mouse | Muller Glia, RPE | 1:1000 | Abcam | Ab15051 | AB_2269474 |
| green fluorescent protein (GFP) | chicken | Donor tissue expressing EGFP | 1:500 | Aves Labs | GFP1010 | AB_2307313 |
| HPC-1 (Syntaxin) | mouse | Synaptic layers, amacrine cells | 1:500 | Dr. Colin Barnstable, Penn State Univ. PA ^1^ | N/A | N/A |
| Iba-1 | rabbit | Microglia | 1:100 -1:200 | Biocare Medical (Pacheco, CA) | CP 290 A | AB_10578940 |
| PKCα (H-7) | mouse | Rod bipolar cells | 1:500 | Santa Cruz | SC-8393 | AB_628142 |
| Recoverin | rabbit | Photoreceptors, cone bipolar cells | 1:500 | Millipore | AB5585 | AB_2253622 |
| Rhodopsin (rho1D4) | mouse | Rods | 1:100 | Dr. Robert Molday, Univ. of British Columbia ^2^ | N/A | N/A |
| Secretagogin (D4V1Y) | rabbit | Cone bipolar cells | 1:200 | Cell Signaling Technology | 14037S | AB_2798371 |

**Supplemental Table S1: Antibodies used in this study**

| **S1.B. Secondary Antibodies** | | | | | | |
| --- | --- | --- | --- | --- | --- | --- |
| **Conjugate** | **species** | **specific for** | **dilution** | **supplier** | **Catalogue #** | **RRID** |
| Alexa Fluor 488 | Donkey | Rabbit IgG (H+L) | 1:400 | Jackson Immuno Research (West Grove, PA) | 711-545-152 | AB_2313584 |
| Alexa Fluor 488 | Donkey | Chicken IgY | 1:400 | Jackson Immuno Research | 703-545-155 | AB_2340375 |
| Alexa Fluor 488 | Donkey | Goat IgG | 1:400 | Jackson Immuno Research | 705-545-003 | AB_2340428 |
| Rhodamine Red-X | Donkey | Mouse IgG (H+L) | 1:400 | Jackson Immuno Research | 715-295-151 | AB_2340832 |
| Alexa Fluor 647 | Donkey | Rabbit IgG (H+L) | 1:400 | Jackson Immuno Research | 711-605-152 | AB_2492288 |

**References:**

1	Barnstable, C. J., Hofstein, R. & Akagawa, K. A marker of early amacrine cell development in rat retina. Brain Res 352, 286-290, doi:10.1016/0165-3806(85)90116-6 (1985).

2	Molday, R. S. & MacKenzie, D. Monoclonal antibodies to rhodopsin: characterization, cross-reactivity, and application as structural probes. Biochemistry 22, 653-660 (1983).

|  | Rat ID | % Visual | Selective |
| --- | --- | --- | --- |
| 1 | 'RNT110' | 9.6% | * |
| 2 | 'RNT118' | 11.5% | * |
| 3 | 'RNT165' | 48.6% | ** |
| 4 | 'RNT169' | 58.6% | ** |
| 5 | 'Rn2762' | 6.0% | * |
| 6 | 'Rn2778' | 20.0% | * |
| 7 | 'Rn2781' | 14.3% | ** |
| 8 | 'Rn2797' | 36.0% | * |
| 9 | 'Rn2829' | 19.4% | * |
| 10 | Rn2751 | 0.0% | - |

**Table S2. Details of response cells proportions for each transplanted rat.**

* Selective cells observed

** results included in the final analyses
